# *PyFolding*: An open-source software package for graphing, simulation and analysis of the biophysical properties of proteins

**DOI:** 10.1101/191593

**Authors:** Alan R. Lowe, Albert Perez-Riba, Laura S. Itzhaki, Ewan R.G. Main

## Abstract

For many years, curve fitting software has been heavily utilized to fit simple models to various types of biophysical data. Although such software packages are easy to use for simple functions, they are often expensive and present substantial impediments to applying more complex models or for the analysis of large datasets. One field that is relient on such data analysis is the thermodynamics and kinetics of protein folding. Over the past decade, increasingly sophisticated analytical models have been generated, but without simple tools to enable routine analysis. Consequently, users have needed to generate their own tools or otherwise find willing collaborators. Here we present *PyFolding*, a free, open source, and extensible Python framework for graphing, analysis and simulation of the biophysical properties of proteins. To demonstrate the utility of *PyFolding*, we have used it to analyze and model experimental protein folding and thermodynamic data. Examples include: (i) multi-phase kinetic folding fitted to linked equations, (ii) global fitting of multiple datasets and (iii) analysis of repeat protein thermodynamics with Ising model variants. Moreover, we demonstrate how *Pyfolding* is easily extensible to novel functionality beyond applications in protein folding via the addition of new models. Example scripts to perform these and other operations are supplied with the software, and we encourage users to contribute notebooks and models to create a community resource. Finally, we show that *PyFolding* can be used in conjunction with Jupyter notebooks as an easy way to share methods and analysis for publication and amongst research teams.

## Introduction

The last decade has seen a shift in the analysis of experimental protein folding and thermodynamic stability data from the fitting of individual datasets using simple models to increasingly complex models using global optimization over multiple large datasets [examples include Refs: (3-21)]. This shift in focus has required moving from user-friendly, but expensive software packages to bespoke solutions developed in computing environments such as MATLAB and Mathematica or by using in-house solutions [examples include: (3, 6, 12, 21, 22)]. However, as these methods of analysis have become more essential, simple curve fitting software no longer provides sufficient flexibility to implement the models. Thus, there is increasingly a need for substantially more computational expertise than previously required. In this respect the protein folding field contrasts with other fields, for example x-ray crystallography, where free or inexpensive and user-friendly interfaces and analysis packages have been developed (23).

Here we present *PyFolding*, a free, open-source and extensible framework for graphing, analysis and simulation. At present, it is customised for the analysis and modelling of protein folding kinetics and thermodynamic stability. To demonstrate these and other functions we present a number of examples as Jupyter notebooks. The software, coupled with the supplied models / Jupyter (iPython) notebooks, can be used by researchers with less programming expertise to access more complex models/analyses and share their work with others. Moreover, *PyFolding* also enables researchers to automate the time-consuming process of combinatorial calculations, fitting data to multiple models or multiple models to specific data. This enables novice users to simply replace the filenames of the datasets with their own and execute the same calculations for their systems. For more advanced users, new models and functionality can be added with ease by utilising the template models. The Jupyter notebooks provided also show how *PyFolding* provides an easy way to share analysis for publication and amongst research teams.

## Results & Discussion

*PyFolding* is distributed as a lightweight, open-source Python library through *github* and can be downloaded with instructions for installation from the authors’ site^1^. *PyFolding* has several dependencies, requiring Numpy, Scipy and Matplotlib. These are now conveniently packaged in several Python frameworks, enabling easy installation of *PyFolding* even for those who have never used Python before (described in the “SETUP.md” file of *PyFolding* and as a series of instructional videos to demonstrate the installation and use of *PyFolding*^2^). As part of *PyFolding*, we have provided many commonly used folding models, such as two‐ and three-state equilibrium folding and various equivalent kinetic variations, as standard (S.I. Jupyter notebook 1-4 & 8). Functions and models themselves are open source and are thus available for inspection or modification by both reviewers and authors. Moreover, due to the open source nature, users can introduce new functionality by adding new models into the library building upon the template classes provided. We encourage users to contribute notebooks and models to create a community resource.

### Fitting and evaluation of typical folding models within *PyFolding*

*PyFolding* uses a hierarchical representation of data internally. Proteins exist as objects that can have metadata as well as multiple sets of kinetic and thermodynamic data associated with them. Input data such as chevron plots or equilibrium denaturation curves can be supplied as comma separated value files (.CSV). Once loaded, each dataset is represented in *PyFolding* as an object, associating the data with numerous common calculations. Models are represented as functions that can be associated with the data objects you wish to fit. As such, datasets can have multiple models and *vice versa* enabling automated fitting and evaluation (S.I. Jupyter notebooks 1-3). Parameter estimation for simple (non-Ising) models is performed using the Levenberg–Marquardt non-linear least-mean-squares optimization algorithm to optimize the appropriate objective function [as implemented in SciPy (24)]. The output variables (with standard error) and fit of the model to the dataset (with R^2^ coefficient of determination & 95% confidence levels) can be viewed within *PyFolding* and/or the fit function and parameters written out as a CSV file for plotting in your software of choice (S.I Jupyter notebook 1-3). Importantly, by representing proteins as objects, containing both kinetic and equilibrium datasets, *PyFolding* enables users to perform and automate higher-level calculations such as Phi-value analysis (25, 26), which can be tedious and time-consuming to perform otherwise (S.I Jupyter notebook 3). Moreover, users can define their own calculations so that more complex data analysis can be performed. For example, multiple kinetic phases of a chevron plot (fast and slow rate constants of folding) can be fitted to two linked equations describing the slow and fast phases of a 3-state folding regime (Figure 1, S.I Jupyter Notebook 4). We believe that this type of fitting is extremely difficult to achieve with the commercial curve fitting software commonly employed for analysing these data, owing to the complexity of parameter sharing amongst different models and datasets.

**Figure 1:**
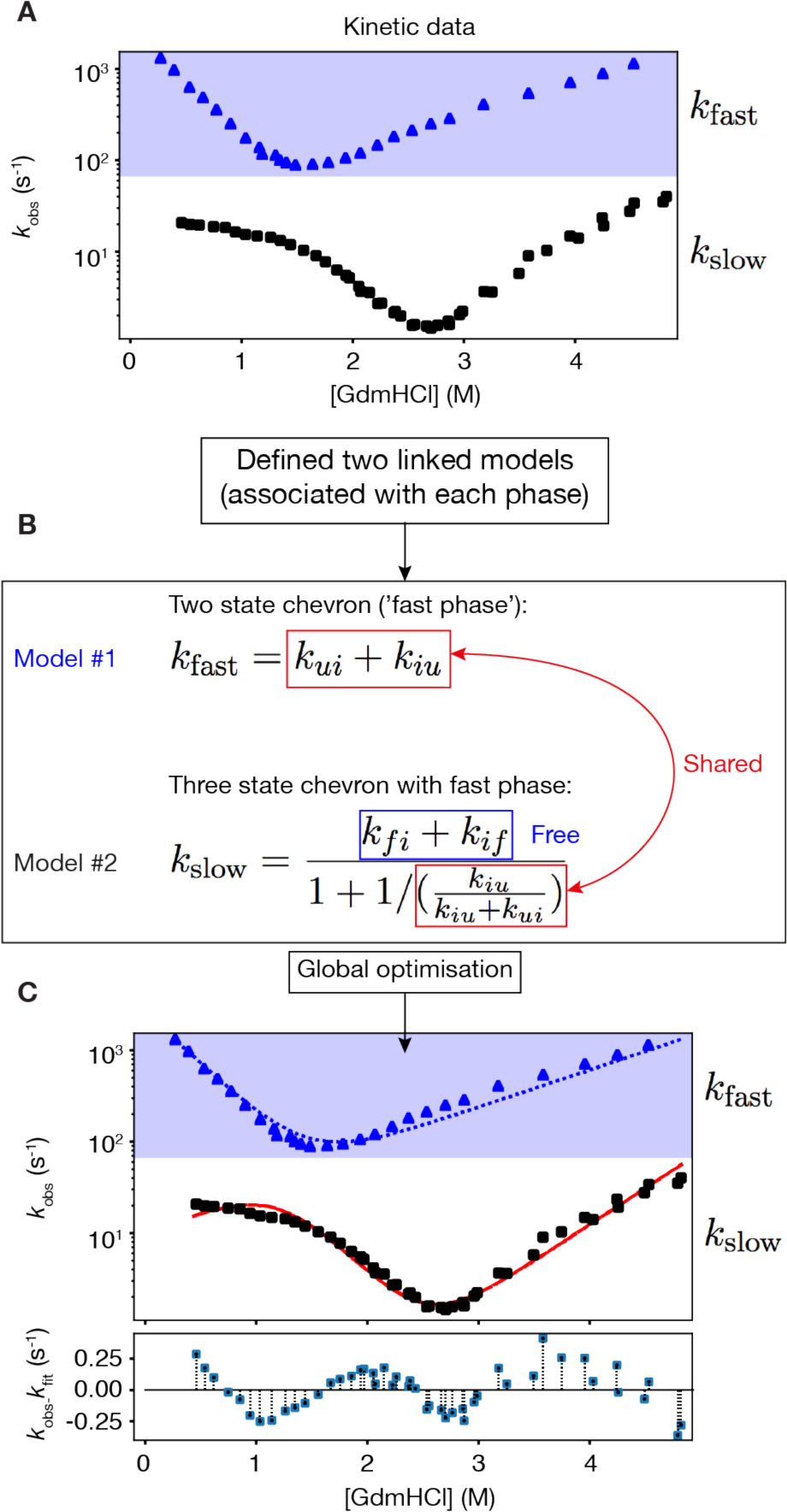
Work flow example of the fitting linked equations in *PyFolding*. **(A)** Unfolding and folding kinetics (chevron plots) showing the distinct fast and slow phases for the 3-state folding thermophilic AR protein (tANK) identified in the archaeon *Thermoplasma*(2) are loaded into *PyFolding* as Chevron objects. **(B)** Two linked models (functions) are associated with the chevron data. These describe the fast (Model #1) and slow phases (Model #2) of the chevrons. Certain rate constants and their associated m-values, are shared between the two models. The other parameters are “free” and associated and fitted only in the slow phase model. **(C)** Global optimization within *PyFolding* enables simultaneous fitting of the two models with shared parameters to the two respective phases. The resultant fits for the fast (blue dotted line) and slow phases (red solid line) are shown overlaid on the observed data. The residuals show the difference between the slow phase observations and fit. These calculations can be found in SI Jupyter Notebook 4.

### More “complex” fitting, evaluation and simulations using the Ising Model

Ising models are statistical thermodynamic nearest-neighbour models that were initially developed for ferromagnetism (27, 28). Subsequently, they have been used with great success in both biological and non-biological systems to describe order-disorder transitions (12). Within the field of protein folding and design they have been used in a number of instances to model phenomena such as helix to coil transitions, beta-hairpin formation, prediction of protein folding rates/thermodynamics and with regards to the postulation of downhill folding (6, 12, 20, 29-34). Most recently two types of one-dimensional (1-D) variants have been used to probe the equilibrium and kinetic un/folding of repeat proteins (3, 12, 17, 21, 22, 35, 36). The most commonly used, and mathematically less complex, has been the 1-D homopolymer model (also called a homozipper). Here, each arrayed element of a protein is treated as an identical, equivalent independently folding unit, with interactions between units via their interfaces. Analytical partition functions describing the statistical properties of this system can be written. By globally fitting this model to, for example, chemical denaturation curves for a series of proteins that differ only by their number of identical units, the intrinsic energy of a repeated unit and the interaction energy between the folded units can be delineated. However, this simplified model cannot describe the majority of naturally occurring proteins where subunits differ in their stabilities, and varying topologies and/or non-canonical interfaces exist. In these cases, a more sophisticated and mathematically more complex heteropolymer Ising model must be used. Here the partition functions required to fit the data are dependent on the topology of interacting units and thus are unique for each analysis.

At present, there is no freely available software that can globally fit multiple folding datasets to a heteropolymer Ising model, and only a few that can adequately implement a homopolymer Ising model. Therefore, most research groups have had to develop bespoke solutions to enable analysis of their data (3, 21, 22, 35, 36). Significantly, in *PyFolding* we have implemented methods to enable users to easily fit datasets of proteins with different topologies to both the homozipper and heteropolymer Ising models. To achieve this goal *PyFolding* presents a flexible framework for defining any non-degenerate 1-D protein topology using a series of primitive protein folding “domains/modules” (Figure 2). Users define their proteins’ 1-D topology from these domains (S.I. Jupyter notebook 5-6). *PyFolding* will then automatically calculate the correct partition function for the defined topology, using the matrix formulation of the model [as previously described (12)], and globally fit the equations to the data as required (S.I. Jupyter notebook 5-6). The same framework also enables users to simulate the effect of changing the topology, a feature that is of great interest to those engaged in rational protein design (S.I. Jupyter notebook 7).

**Figure 2:**
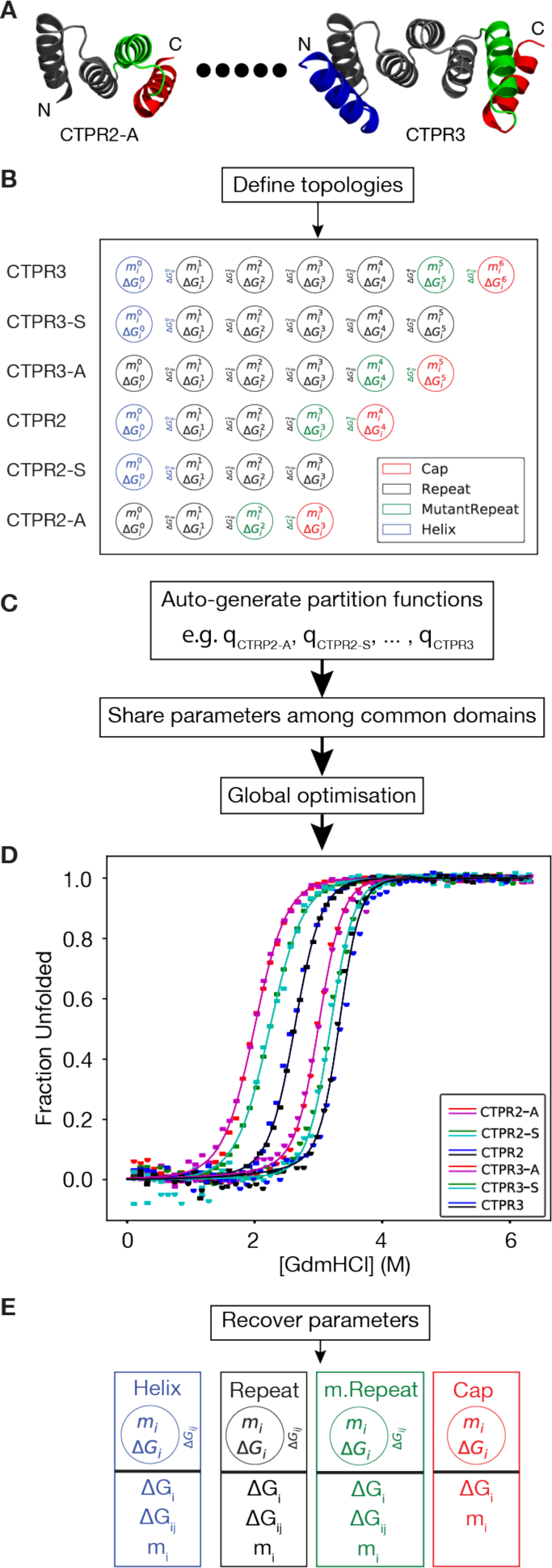
Work flow example of global optimization of a Heteropolymer Ising model in *PyFolding*. **(A)** GdmHCl-induced equilibrium denaturations of a series of single-helix deletion CTPRn proteins are loaded into *PyFolding* as EquilibriumDenaturation objects. In the figure we schematically represent these as individual protein structures corresponding to the smallest in the series (CTPR2-A) upto (dots) the largest (CTPR3) (3). The figures were made with Pymol and individual helices are coloured by the user defined topology used by the ising model - Helix (blue), Repeat (black), a mutant Repeat (green) or a Cap (red). **(B)** Using *PyFolding*’s built-in primitive protein folding “domains/modules”, one can define topologies for each protein in the series. Each primitive is a container for several thermodynamic parameters to describe the intrinsic and interfacial stability terms. **(C)** Using the topologies defined in (B), *PyFolding* will automatically generate the appropriate partition functions (q) for each protein in the series using a matrix formulation, and share parameters between other proteins in the series. **(D)** A final global fitting step finds the optimal set of parameters to describe the series. **(E)** The optimal parameters (and their estimated errors/confidence intervals) for each domain primitive are recovered and output for the user. These calculations can be found in SI Jupyter Notebook 6.

To determine a globally optimal set of parameters that minimises the difference between the experimental datasets and the simulated unfolding curves, *PyFolding* uses the stochastic differential evolution optimization algorithm (37) implemented in SciPy (24). In practice, experimental datasets may not adequately constrain parameters during optimization of the objective function, despite yielding an adequate curve fit to the data. It is therefore essential to carefully assess the output of the model to verify the validity of any topologies and the resultant parameters. A description of how *PyFolding* provides the error estimates and determines how constrained parameters are is given in the error analysis section below. As with the simpler models, *PyFolding* can be used to visualise the global minimum output variables (with standard errors) and the fit of the model to the dataset (with R^2^ coeff. of determination) (S.I. Jupyter notebook 5-6). The output can also be exported as a CSV file for plotting in your software of choice. In addition, *PyFolding* outputs a graphical representation of the topology used to fit the data and a graph of the denaturant dependence of each subunit used (Figure 2). Thus, *PyFolding* enables non-experts to create and analyse protein folding datasets with either a homopolymer or heteropolymer Ising model for any reasonable 1-D protein topology. Moreover, once the 1-D topology of your protein has been defined, *PyFolding* can also be used to simulate and thereby predict folding behavior of both the whole protein and the sub-units that it has been composed of (S.I. Jupyter notebook 7). In principle, this type of approach could be extended to higher dimensional topologies, thus providing a framework to enable rational protein design.

### Error Analysis

We calculate various metrics to assess the quality of the output from *PyFolding*. All independent non-constant variables are reported with a standard error of each parameter, *i*:

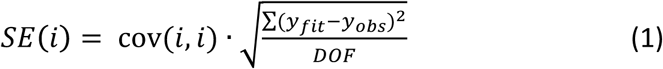

Where cov is the covariance matrix (where cov(*i, i*) represents the variance of parameter *i*), *y_fit_* are the y-values of the fit at the observed x values, *y_obs_* are the observed y values of the data and *DOF* represents the degrees of freedom (the number of data-points minus the number of free variables). From these values we can also calculate the confidence interval (nominally at 95%) where, the confidence interval for parameter *i* is:

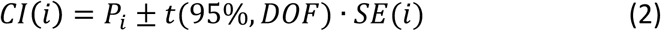

Where *P_i_* is the value of parameter *i* and *t*(95%, *DOF*) is the t-distribution at 95% with *DOF* degrees of freedom. Finally, we report the coefficient of determination (*R^2^*) as a statistical measure of the error between the data and the fitted model:

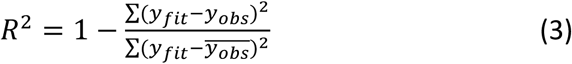

Where 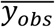 represents the mean of the observed data.

In all models other than the heteropolymer Ising model we utilise a gradient optimiser such as the Levenberg-Marquardt algorithm that yields a covariance matrix of the fitted parameters. However, since we must utilise a different optimization method (the differential evolution optimiser) for the global fitting of heteropolymer Ising models, we calculate the errors in a slightly different way. The optimiser does not yield a covariance matrix as default, so we calculate a numerical approximation based on the Jacobian matrix (here, a matrix of numerical approximations of all the partial differentials of all variables) as follows:

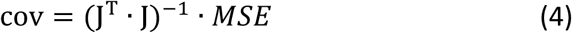

where **J** is the Jacobian matrix, and *MSE* is the mean squared error of the fit.

In *PyFolding* we have provided estimates of the standard error and confidence intervals for each parameter (calculated as described above) using this numerical approximation of the covariance matrix. In general, estimating errors for the parameters or the uniqueness of the solution in heteropolymer models is a complex problem, owing to the method of optimization used. Interestingly, Barrick and coworkers used Bootstrap analysis to evaluate parameter confidence intervals (12). However, many of the published studies either do not describe how error margins were determined or simply list the error between the data and curve fit. Here, when confronted with ill-posed datasets or poorly chosen topologies, which can produce an adequate curve fit to the data (as measured by *R^2^*), *PyFolding*’s numerical error approximation becomes unstable leading to large errors. Thus, in evaluating the determinant of the Jacobian as well as the estimated errors it is possible to assess the quality of the model and the validity of the solution - large errors show that the model parameters are not properly constrained. In such cases, *PyFolding* raises the appropriate warnings to enable the user to quickly interpret the results and adjust the topologies and members of a dataset appropriately.

## Conclusion

Here we have shown that *PyFolding*, in conjunction with Jupyter notebooks, enables researchers with minimal programming expertise the ability to fit both “typical” and complex models to their thermodynamic and kinetic protein folding data. The software is free and can be used to both analyse and simulate data with models and analyses that expensive commercial user-friendly options cannot. In particular, we have incorporated the ability to fit and simulate equilibrium unfolding experiments with user defined protein topologies, using a matrix formulation of the 1-D heteropolymer Ising model. This aspect of *PyFolding* will be of particular interest to groups working on protein folds composed of repetitive motifs such as Ankyrin repeats and TPRs, given that these proteins are increasingly being used as novel antibody therapeutics (38-41) and biomaterials (42-47). Further, as analysis can be performed in Jupyter notebooks, it enables novice researchers to easily use the software and for groups to share data and methods. We have provided a number of example notebooks and accompanying video tutorials as a resource accompanying this manuscript, enabling other users to recreate our data analysis and modify parameters. Finally, due to *PyFolding*’s extensible framework, it is straightforward to extend, thus enabling fitting and modelling of other systems or phenomena such as protein-protein and other protein-binding interactions. Such extensions can be rapidly and seamlessly deployed as a community resource thus broadening the functionality of the software.

## Acknowledgements

We would like to thank Dr. Jonathan Phillips for insightful discussion, helpful comments and suggestions. LSI acknowledges the support of a Senior Fellowship from the UK Medical Research Foundation. AP was supported by a BBSRC Doctoral Training Programme scholarship and an Oliver Gatty Studentship. ERGM and LSI labs acknowledge support from a Leverhulme Trust project grant.

## Footnotes

1 https://github.com/quantumjot/PyFolding

2 https://github.com/quantumjot/PyFolding/wiki

